# X-ray microscopy enables multiscale high-resolution 3D imaging of plant cells, tissues, and organs

**DOI:** 10.1101/2020.12.18.423480

**Authors:** Keith E. Duncan, Kirk J. Czymmek, Ni Jiang, August C. Thies, Christopher N. Topp

**Affiliations:** Donald Danforth Plant Science Center, Saint Louis, Missouri, 63132, USA

## Abstract

Capturing complete internal anatomies of plant organs and tissues within their relevant morphological context remains a key challenge in plant science. While plant growth and development are inherently multiscale, conventional light, fluorescence, and electron microscopy platforms are typically limited to imaging of plant microstructure from small flat samples that lack direct spatial context to, and represent only a small portion of, the relevant plant macrostructures. We demonstrate technical advances with a lab-based X-ray microscope (XRM) that bridge the imaging gap by providing multiscale high-resolution 3D volumes of intact plant samples from the cell to whole plant level. Serial imaging of a single sample is shown to provide sub-micron 3D volumes co-registered with lower magnification scans for explicit contextual reference. High quality 3D volume data from our enhanced methods facilitate more sophisticated and effective computational segmentation and analyses than have previously been employed for X-ray based imaging. Advances in sample preparation make multimodal correlative imaging workflows possible, where a single resin-embedded plant sample is scanned via XRM to generate a 3D cell-level map, and then used to identify and zoom in on sub-cellular regions of interest for high resolution scanning electron microscopy. In total, we present the methodologies for use of XRM in the multiscale and multimodal analysis of 3D plant features using numerous economically and scientifically important plant systems.

## INTRODUCTION

Visible features of the plant, such as flowers, leaves and roots, result from microscopic processes of tissue and organ formation, which are in turn driven by cellular and molecular dynamics. How whole plants and their organs, tissues, cells, organelles, and sub-cellular molecules interact within three-dimensional (3D) space is crucial to understanding plant biology across diverse genetic backgrounds and changing environments. Capturing 3D external and internal volume data brings an important spatial component to plant phenotype analysis that can help identify genetic factors that control trait development not manifested in simpler 2D images. Recently, 3D imaging platforms and corresponding phenotyping methods have developed rapidly and many approaches have been applied to a wide diversity of plant growth and development phenomena (Morris et al., 2017; Strock et al., 2019; Jiang et al., 2019; Tracy et al., 2017; Roth et al., 2019; Li et al., 2020; Perez-Sanz et al., 2017; Czymmek et al., 2020). Furthermore, sophisticated analysis pipelines have been developed to extract and quantify data from relevant features, with improved imaging leading directly to improved trait analysis and models (Wolny et al., 2020; Sultan et al., 2020; Théroux-Rancourt et al., 2020; McKay Fletcher et al., 2020; Jackson et al., 2019; Bassel and Smith, 2016; Koebernick et al., 2017; Mathers et al., 2018).

Current 3D systems use photons, electrons, and X-rays for imaging biological samples. Photon-based tomography (confocal, multiphoton, lightsheet, super-resolution) generates valuable 3D information with chemical specificity based on fluorescent probes and/or auto-fluorescence in both living and fixed samples (Wymer et al., 1999; Feijó and Moreno, 2004; Schubert, 2017; Ovečka et al., 2018). However, imaging with photons fundamentally has sample size constraints (Prunet and Duncan, 2020) and non-destructive approaches (but see Strock et al., 2019) are typically limited to shallow image depths for plant tissues as they are highly refractive due to cell walls, vacuoles, and air spaces, that often require special preparation or clearing to mitigate (Littlejohn et al., 2010; Kurihara et al., 2015). Another approach, 3D volume scanning electron microscopy (SEM), repeatedly images a resin block-face after removing thin layers via an *in situ* microtome or surface ablation via focused ion beam (Bhawana et al., 2014; Kittelmann et al., 2016; Czymmek et al., 2020; Harwood et al., 2020). With volume SEM, detailed 3D spatial information is possible down to a few nanometers, however practical imaging volumes are typically restricted to a maximum of a few hundred micrometers (Kubota et al., 2018). Additionally, sample preparation is highly specialized, where prolonged protocols with greatly increased sample staining are required to ensure adequate signal-to-noise and sample resolution.

The X-ray microscope (XRM) is a specialized X-ray tomography instrument that incorporates microscope objectives. In XRM, 2D digital radiographs are projected onto a scintillator-coated objective lens where the X-ray signal is converted into visible light, magnified, and the resulting image collected with a detector, typically a CCD camera. The objective lens in the beam path increases the resolution of the XRM compared to conventional X-ray tomography systems, which rely solely on geometric magnification, and multiple objective lenses make possible the collection of 3D volumes over a wide range of sample sizes and resolutions not practical with other imaging platforms. Furthermore, in contrast to photon- and electron-based tomography, where imaging penetration depths are restricted to one or a few cell layers, the ability of X-rays to penetrate through large, intact and highly light scattering plant structures is a significant advantage. Thus, lab-based XRM provides a critical link between photon and electron 3D imaging, establishing high-resolution intermediate magnification levels for whole plant imaging.

X-ray imaging contrast is based on differential density of features within a specimen, however most biological organisms are composed of relatively uniform low-density organic material, often with insufficient contrast to generate useful 3D image data. For this reason, sample fixation protocols and contrast agents play important roles in imaging biological samples with XRM (Metscher, 2009). Staedler *et al.* (2013) evaluated conventional EM sample fixation methods and contrast agents as tools for examining plant samples with XRM. These methods were sufficient for the relatively short scan times necessary to capture floral anatomy for analysis using basic grayscale thresholding, but do not provide adequate stability for the longer scan times required for high magnification/resolution of more subtle and complicated plant features.

Here, we have built upon these earlier methods, incorporated improvements in sample preparation, and leveraged instrument technological advances to expand the range of plant cells, tissues, and organs that can be imaged at high resolution with XRM. This enabled unprecedented 3D perspectives and insights for phenotype and form-function studies. Further, due to the improved image quality of 3D volume data, advanced image analysis and cell-level segmentation of relevant structures of interest using computer vision and learning is significantly enhanced. We applied XRM with improved sample preparation for multimodal and multiscale 3D imaging, where samples prepared for high-resolution volume SEM were first imaged with XRM as part of a correlative workflow (Bushong et al., 2015; Tsang et al., 2018; Mitchell et al., 2019; Bayguinov et al., 2020). Taken together, we anticipate adoption of these improved techniques will make a significant contribution to plant biology, expanding the reach of XRM as a routine tool for 3D imaging for plant scientists.

## RESULTS^1^

### Meristem Biology

Our ultimate goal was to develop plant specific XRM protocols and workflows that improved specimen staining and contrast with excellent stability for long duration high-resolution scans. To image plant tissue with XRM, differential contrast of cell walls was particularly useful, however this is difficult to achieve, especially in cytoplasm-rich meristematic tissue. For wet preparations, we routinely achieved high contrast using ethanolic phosphotungstic acid (ePTA) and improved sample stability by mounting specimens in low melting point (LMP) agarose (see Supplemental Diagram 1). Longer scans were made possible by improved sample contrast and stability, producing better signal-to-noise ratios and edge sharpness in images, both of which helped differentiate individual meristematic cells within inflorescence structures. Figure 1 illustrates how our XRM approach addressed this challenge with examples of meristematic tissue from *Setaria viridis* and *Zea mays*. Figures 1A,1E show ePTA contrasted shoot apical meristems from *S. viridis* and *Z. mays*, respectively. These methods were also effective for generating detailed 3D volumes of inflorescence structures that allowed comprehensive visualization and segmentation of spikelet pairs and sterile bristles in *S. viridis* (Figures 1B,C,D) where the inherent difference in tissue density between spikelets and bristles was observed as variations in grayscale. Similar imaging detail was achieved in ear primordia of *Z. mays* (Figures 1F,G) where inflorescence, spikelet pair, spikelet, and floral meristem features were well-contrasted and resolved for 3D measurement.

**Figure 1.**
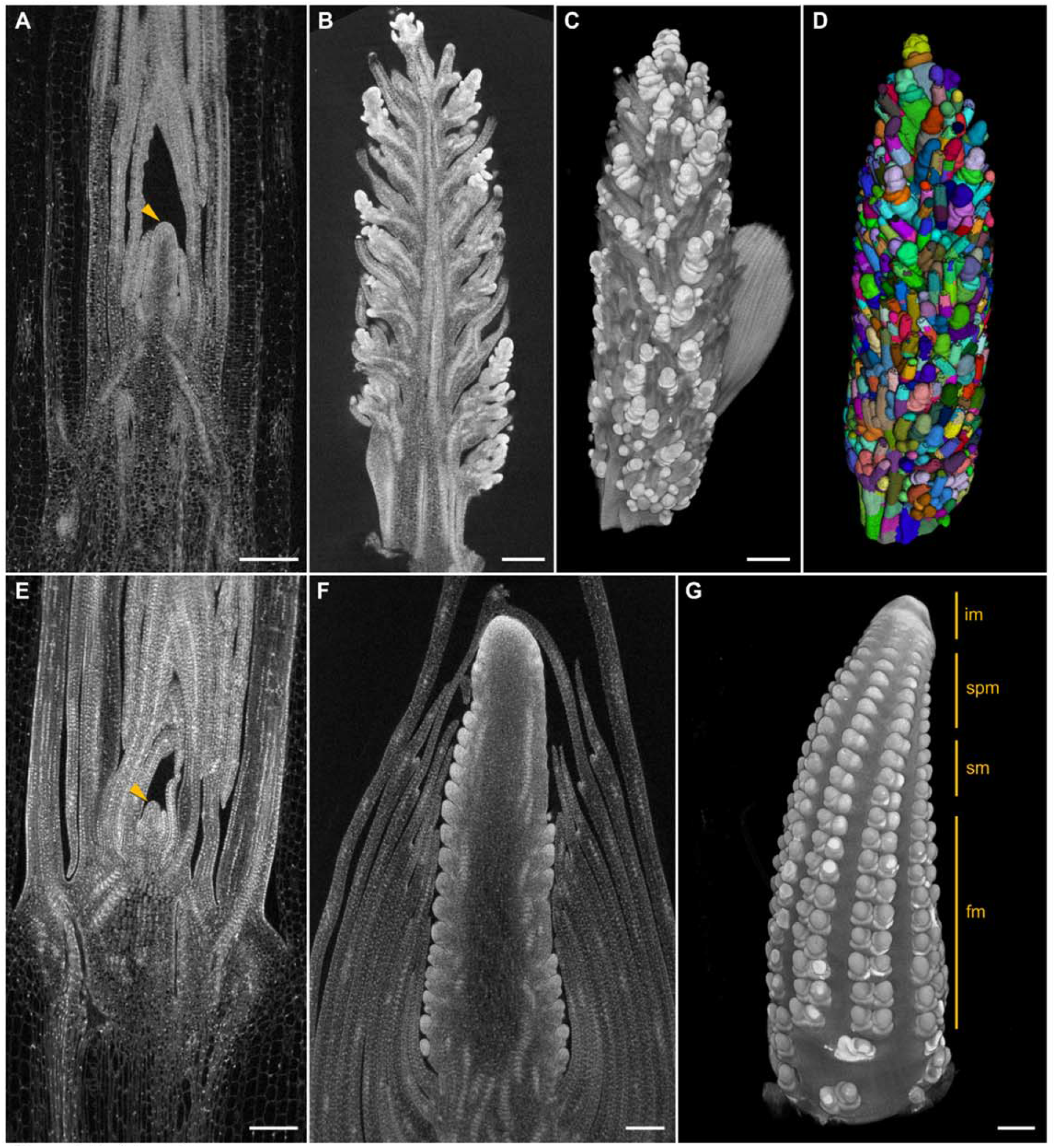
Meristem biology Shoot apical meristems (**arrowheads**) from *Setaria viridis* (**A**) and *Zea mays* (**E**) illustrate how development of this important organ can be visualized in 3D at high resolution while remaining intact within surrounding sheath tissue. Inflorescence structure from *S. viridis* was scanned (**B**), displayed as a volume rendering (**C**), and segmented (**D**) to identify individual spikelets and bristles. X-ray microscope imaging of *Z. mays* ear primordium (**F**), with the volume rendering (**G**) providing high resolution 3D details of regions containing developing inflorescence (**im**), spikelet pair (**spm**), spikelet (**sm**), and floral meristems (**fm**). All scale bars 200 μm.

### Floral anatomy and development

Next, we applied this approach to soybean (*Glycine max*), whose indeterminate growth regularly produces flowers at multiple developmental stages. The relatively large (~1 cm) axillary bud complex (Figure 2A inset) was difficult to image as a single unit in 3D with other microscopy technologies, but was imaged in its entirety with XRM as shown in Figure 2A. Reproductive structures including anthers, pollen, ovules, and the stigmatic surfaces from multiple developmental stages can be seen in high detail (Figure 2A). A higher resolution scan of a single developing flower (Figure 2B) shows the 3D spatial relationships of pollen-filled anthers to the stigma, stigmatic surface, and ovules. Paired ovule scans demonstrate the multiscale capability of XRM (Figures 2C,D). An overview scan at 1.5 μm voxel resolution of two pollinated ovules (Figure 2C) was used to select a specific ovule-of-interest for a higher resolution scan at 0.6 μm voxel resolution (Figure 2D), where details like the egg cell nucleus, fused polar nuclei, and developing synergids were evident. This example clearly shows the effectiveness of these methods for generating high resolution scans over different scales from a single mounted specimen. Additional examples of the benefits of multiscale imaging with XRM include *S. viridis* and *Eragrostis tef* (teff, Supplementary Figure 1) and *Thlaspi arvense* (pennycress, Supplementary Figures 2C,D), where multiscale 3D imaging of single sample captured entire intact inflorescence structures, followed by nanometer scale imaging of internal structures such as cotyledons, ovaries, ovules, stigmas, and anthers. Multiscale imaging of *A. thaliana* plantlets also demonstrated the suitability for visualization of the shoot apical meristem in agarose-stabilized samples (Supplementary Figures 2A,B).

**Figure 2.**
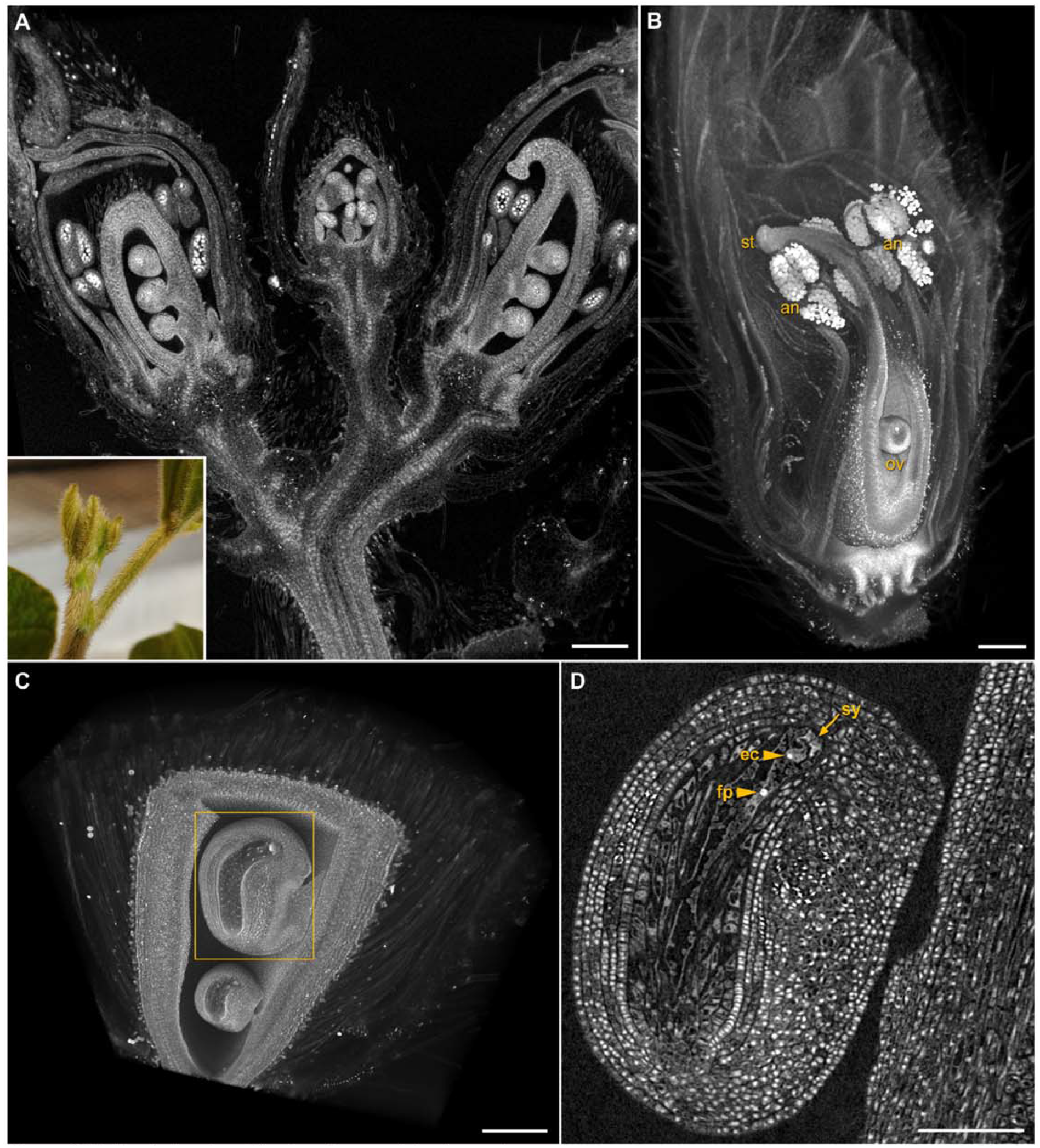
Multiscale imaging of floral anatomy and development Fixed and contrast enhanced reproductive structures of *Glycine max*. Axillary buds (**A,inset**) contrasted in phosphotungstic acid, the entire intact sample imaged with XRM for a detailed 3D volume of this complicated structure; scale bar 300 μm. **B** Volume rendering of a single developing flower shows the relationship of pollen-filled anthers (**an**), ovules (**ov**), and stigmatic surface (**st**) to one another in 3D space; scale bar 200 μm. **C** Volume rendering of developing ovules; scale bar 200 μm. **D** High magnification scan of indicated ovule from **C** illustrates cell-level resolution of egg cell (**ec**) and fused polar (**fp**) nuclei, synergids (**sy**); scale bar 100 μm.

### Root biology

Extension and curvature of roots in 3D space is driven by cell-division and elongation patterns, but the roots of most plants are too thick for light microscopy to capture internal tissue layers that are critical to these dynamics. XRM scanning of fixed and contrast enhanced *Z. mays* root tips (Figures 3A,D) allowed computational segmentation at both the cell (Figure 3B) and tissue level (Figure 3C). Here, root tips were contrasted with either aldehyde-osmium protocol (Methods, Figure 3A) or in ePTA (Methods, Figure 3D), and stabilized in agarose for the long scan times required to visualize individual cell walls and nuclei throughout the full 3D volume (Figure 3D). In a thick sample of a maize node (5 mm), vascular tissues showed excellent contrast with iodine-based staining (Figure 3E) and the complicated architecture of xylem, phloem, and brace root primordia at the nodal plexus were readily observed.

**Figure 3.**
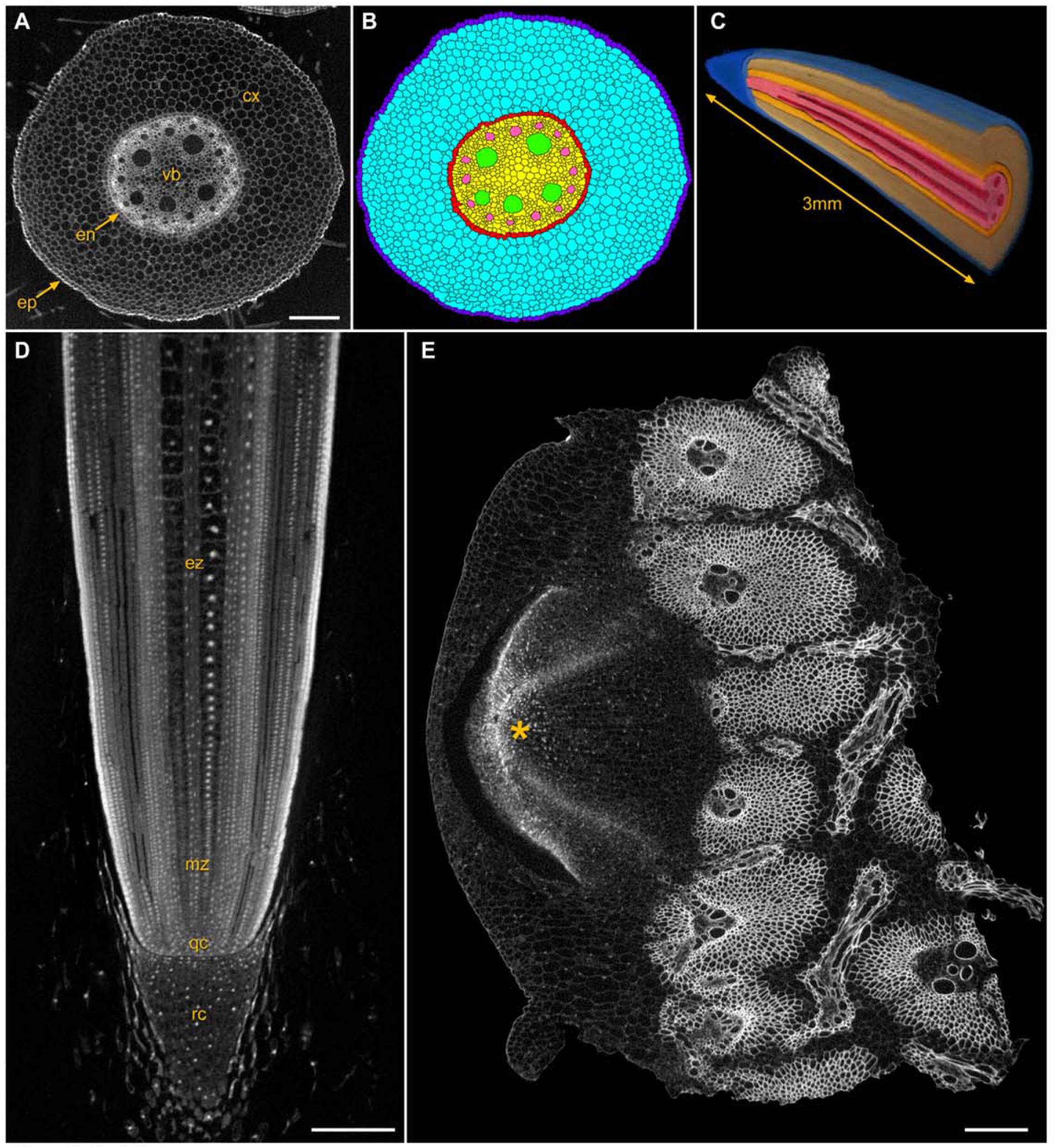
Root biology **A-C***Zea mays* root tip fixed and contrast enhanced to visualize distinct cell layers: epidermis (**ep**), cortex (**cx**), endodermis (**en**), vascular bundle (**vb**), with computational segmentation at both the cell (**B**) and tissue (**C**) levels; scale bar 200 μm. **D** Longitudinal view through the root tip where nuclei are easily visualized in the root cap (**rc**), quiescent center (**qc**), and meristematic (**mz**) and elongation (**ez**) zones; scale bar 200 μm. **E** Stalk sample from the second above-ground node of a *Z. mays* plant at V8, contrasted in Lugol’s iodine, showing a brace root primordium (**\***) and the well-contrasted vascular bundles; scale bar 300 μm.

### Plant-microbe interactions

3D visualization of plant-microbe relationships *in situ* and across scales can provide valuable data about how organisms interact in response to various environmental stimuli and genetic backgrounds. For example, nitrogen-fixing bacteria colonize roots of many leguminous crops, providing host-accessible forms of nitrogen to the plant in exchange for carbon-based energy and a protected environment (Coba de la Peña et al., 2017). Here, *Bradyrhizobium japonicum* was applied to wild-type *G. max* seedlings and the resulting nodules harvested after 14 days. In Figure 4A a root segment with four nodules was imaged with a two-panel stitched XRM scan, and a subsequent higher resolution scan in a selected region of the nodule (Figure 4B) showed well-contrasted plant cells and their nuclei, densely packed with symbiosomes and the surrounding nodule vascular network. The strong affinity of PTA for nuclei (Hayat, 1993) assisted with segmentation, providing a consistent central feature to guide identification of individual cells, combined with the sharp edge contrast of the surrounding plant cell walls allowed detailed segmentation via a combination of manual and automatic approaches (Figures 4C,D).

**Figure 4.**
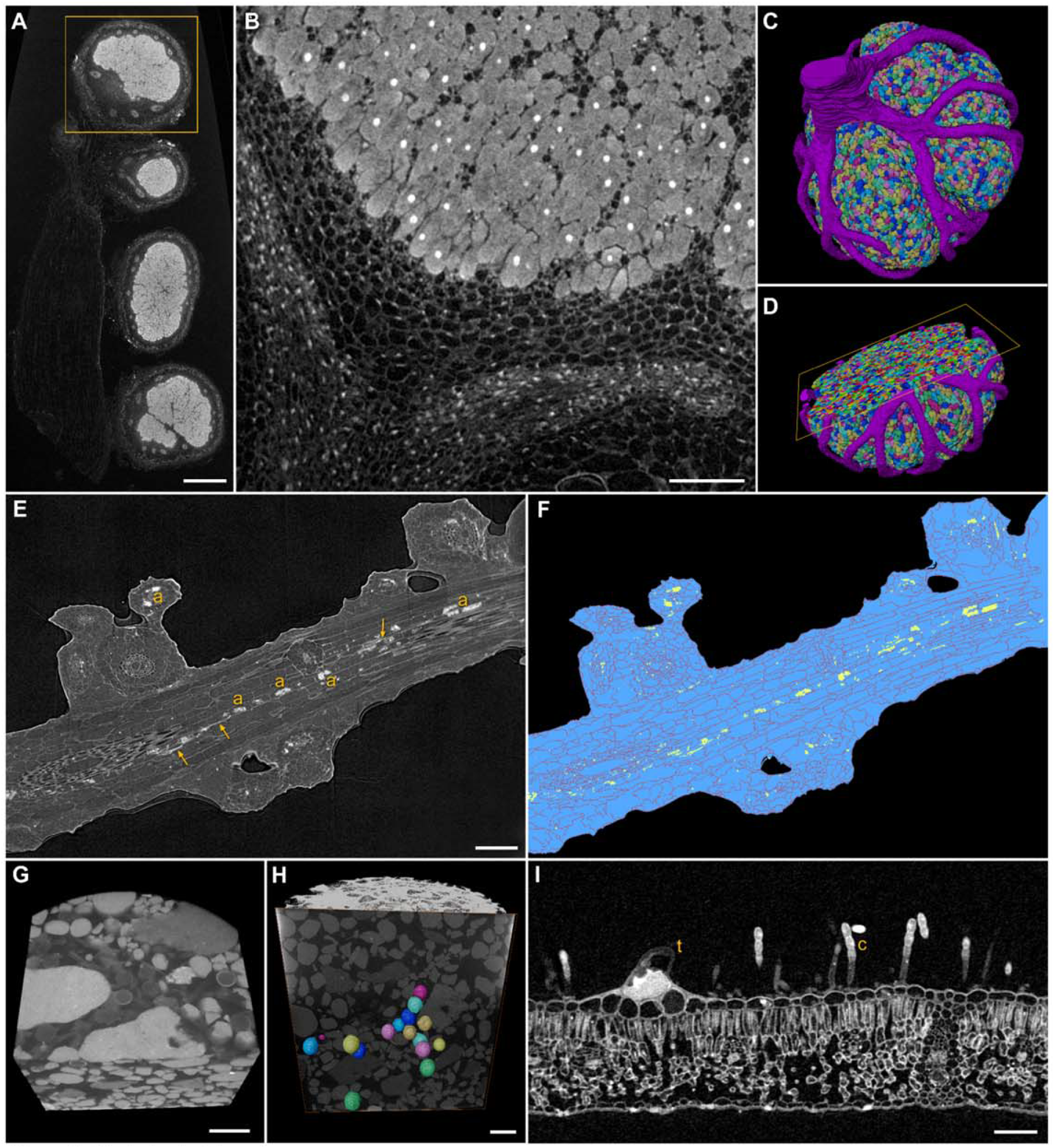
Plant-microbe interactions **A** Low magnification scan of *Bradyrhizobium japonicum* nodules on soybean root; scale bar 500 μm. **B-D** High resolution scan and segmentation of nodule from **A**. Well-contrasted nuclei and cell walls allow segmentation of symbiosomes and vasculature; scale bar 150 μm. **E,F** Colonization of maize root by *Rhizophagus irregularis*, arbuscules (**a**) and intercellular hyphae (**arrows**) segmented in yellow; scale bar 200 μm. **G,H** Volume rendering of *Gigaspora margarita* spores *in situ* scan with spores segmented in color; scale bars 500 μm. **I** Trichome (**t**) and fungal conidiophores with conidia (**c**) on *Echinacea* leaf surface; scale bar 150 μm.

Mycorrhizal fungi colonize root cortical cells from most plant families, providing water and nutrients to the host in exchange for sugars and fatty acids (Luginbuehl and Oldroyd, 2017; Chen et al., 2018). To test the suitability of our XRM methods for mycorrhizal associations, wild-type *Z. mays* plants were grown in the greenhouse in a mixture of field soil and sand, inoculated with spores of the arbuscular mycorrhizal fungus *Rhizophagus irregularis*, and colonized roots were harvested after six weeks. These roots were fixed and contrast enhanced with ePTA and imaged. As Figure 4E illustrates, arbuscules—the symbiotic structures formed within cortical cells to exchange nutrients between host plant and fungus—were clearly visible within maize cortical cells. Intercellular hyphae were also evident by segmentation of these structures shown in Figure 4F. In an alternative system, we grew *Medicago sativa* in 20 mL syringe barrels with a sand-Agsorb® mixture inoculated with spores of *Gigaspora margarita.* To image both symbiotic partners over scales, we first used the XRM flat panel detector which allowed us to visualize the entire root system in a 10 cm x 2 cm syringe barrel. We then scanned specific regions of interest within the barrel (using the 0.4X and 4X objective lenses), to localize *G. margarita* spores *in situ* (Figures 4G,H, Supplementary Figure 4). This multiscale approach yielded unperturbed 3D volume data of fungal structures in relation to the entire host root system without removing the plant from the soil. To evaluate host-pathogen interactions, we imaged powdery mildew infected leaves of *Echinacea* spp. that were collected from a prairie grassland ecosystem, fixed in buffered aldehydes and contrast enhanced with osmium tetroxide, and imaged via XRM. Conidiophores with conidia emerged from the surface of the leaf in Figure 4I, adjacent to host trichomes.

### Correlative microscopy

One challenge facing plant biologists applying high-resolution electron microscopy is targeting specific or rare structures within the plant and/or locating very small regions acquired by EM in the larger bulk plant tissue. This becomes especially problematic when dealing with opaque heavy metal stained and resin embedded EM plant samples, where features are often not visible in the densely stained and osmicated tissue. Here we applied an enhanced staining approach used to achieve uniform staining for serial block face electron microscopy (SBFEM) in brain tissues (Hua et al., 2015) to *Nicotiana benthamiana* leaf tissue which had been embedded in epoxy resin. Our goal was to provide a high contrast/resolution protocol for tissue and subcellular details that was optimized for multi-scale correlative XRM/SBFEM. With this preparation, we generated an overview 3D XRM tomogram using a 4X objective lens at a 1.2 μm voxel size (Figure 5A). Under these conditions, a 3 mm diameter leaf punch could be readily interrogated and clearly revealed the spatial distribution of plant structures including trichomes, palisade mesophyll, spongy mesophyll and vascular tissue at arbitrary slice perspectives (Figure 5B). Subsequently, an XRM scan using the 40X objective lens was applied at 0.2 μm voxel size for visualization of a high-resolution subset of the leaf that was submerged into the lower resolution dataset (Figures 5A,B green cylinder). A representative 2D slice of the *N. benthamiana* leaf cross-section within the high-resolution volume (Figure 5C) shows spatial distribution and more detailed tissue features, and enabled precise targeting of individual palisade cells for subsequent imaging (Figure 5C yellow box, SBFEM overlay). We next targeted the palisade mesophyll for high-resolution SBFEM in the exact same specimen to directly correlate individual chloroplasts at nanometer scale within the 3D XRM volume (Figures 5D,E). The enlarged single 2D slice (Figure 5D) corresponding to Figure 5C from a series of ~1500 SBFEM images of a palisade cell showed multiscale image correlation and highlights the resolution gain from SBFEM for nanoscale features. Subcellular structures such as starch, thylakoid membranes, plastoglobules and individual grana and stacking were readily documented. With the strong staining provided by this protocol combined with the stability afforded by epoxy resin embedding for long term high-resolution XRM imaging, we were able to delineate individual isolated chloroplasts, and sub-chloroplast voids corresponding to starch granules (Figure 5E), despite the fact that the cytoplasmic cell periphery was often densely packed, with contrast largely coming from intensely stained chloroplast membranes. Segmentation using machine learning of high-resolution leaf features permitted clear 3D visualization and distribution of nuclei, cell walls/vascular bundle, chloroplast distribution and their starch granules across tissues (Figure 5F).

**Figure 5.**
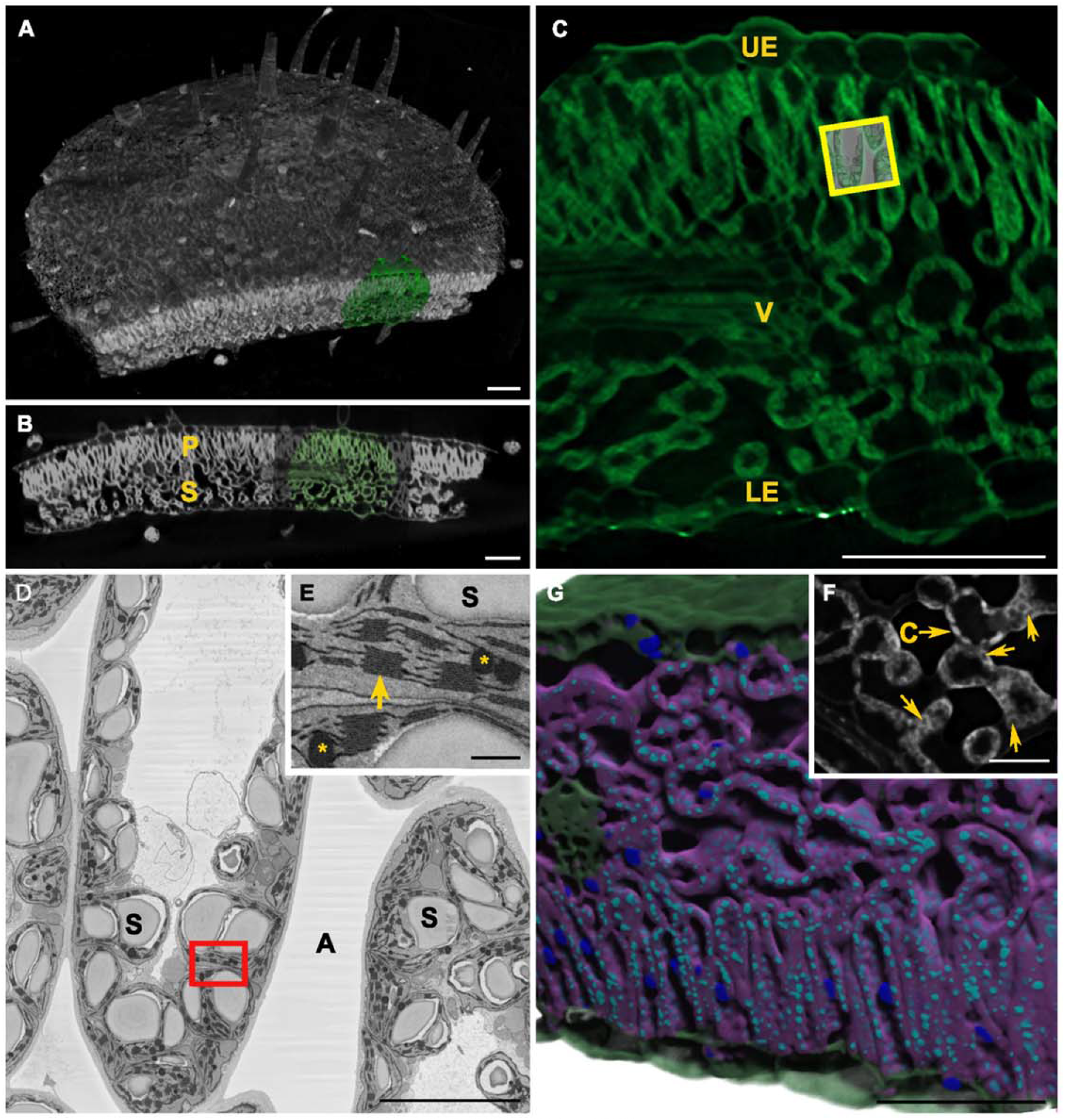
Multiscale correlative X-ray and electron microscopy of tobacco leaf Resin embedded *Nicotiana benthamiana* leaf prepared using OTO staining protocol provided high contrast 3D (**A**) and 2D (**B**) XRM perspectives for corresponding high-resolution XRM (**C**, **A&B green**). This approach provided 3D detail of the palisade (**P**) and spongy (**S**) mesophyll and cell wall defining the vascular tissue (**v**) and upper (**ue**) and lower (**le**) epidermis enabling rich content for subsequent correlative SBFEM (**C EM overlay, yellow box**) at intermediate resolution of palisade cells (**D, corresponding to C yellow box**) with complement of cell organelles, including starch (**s**) and apoplast (**a**) and high-resolution inset (**E, corresponding to D red box**) with grana (**arrow**) and plastoglobules (**asterisks**). High-resolution XRM in spongy mesophyll shows chloroplast (**c**) and starch voids (**arrows**). Deep learning of XRM data allowed 3D segmentation/visualization of nuclei (**dark blue**), starch granules (**light blue**), cell cytoplasm/chloroplasts (**purple**), cell walls (**green**). Scale bars 100 μm (**A-C,G**), 10 μm (**D**), 500 nm (**E**), 20 μm (**F**).

## DISCUSSION

Rapid technological and computational improvements over the past several decades have enabled the acquisition and analysis of 3D volumetric data of ever-increasing size and complexity for plant biology. However, intact plant structures are often difficult to image in 3D with light and/or electron microscopy platforms due to challenges including relatively large sample size, an optically scattering and chemically resistant cuticle, cell walls and air pockets that impede fixation and exogenous treatments, complex morphologies, and delicate features. As a result, a notable gap exists with light and electron imaging modalities that restricts high-resolution tomographic imaging to small samples that provide limited contextual information relative to the whole organ or plant.

Because of its ability to penetrate dense tissues, conventional X-ray computed tomography (X-ray CT) was utilized to address some of these imaging limitations in plant science as early as the 1990s (Aylmore, 1993; Heeraman et al., 1997) and 2000s (Stuppy et al., 2003; Kaminuma et al., 2008; Leroux et al., 2009; Dhondt et al., 2010; van der Niet et al., 2010). However most conventional X-ray CT instruments rely solely on geometric magnification, and thus have functional resolution and sample size limits based on source-sample-detector geometry. Synchrotron-based X-ray CT has advantages in shorter scan times and allows element mapping using X-ray fluorescence (Kopittke et al., 2018), however their limited availability relative to lab-based XRM restricts wide-spread utilization in plant biology. Here, we demonstrated that lab-based XRM could image intact plant structures in fine detail, and subsequently zoom to selected regions of interest down to the cellular and subcellular level, effectively bridging the tomographic imaging gap of light and electron microscopy.

Contrast and stabilization of samples are critical for high-resolution X-ray imaging due to the longer scan times. Previous XRM studies used conventional sample fixation, contrast enhancement, dehydration, and critical point drying followed in some cases by sputter coating (Staedler et al., 2013; Bellaire et al., 2014; Jeiter et al., 2018, 2020). Other researchers stabilized samples in the contrast enhancement medium, sealed within pipette tips or polypropylene tubes (Staedler et al., 2013; Gamisch et al., 2013; Staedler et al., 2018; Reich et al., 2020). In these studies, the goal was to survey a relatively large number of samples and evaluate basic 3D shapes using scans no longer than necessary to achieve stated research aims in a timely manner. With plant tissue prepared in this way, even with floral samples stabilized in polyester fibers or in ethanol, sample drift during longer XRM scans can be problematic and therefore limit achievable resolution and magnification. To address this challenge, we specifically developed a simple but effective strategy whereby fixed samples were contrasted in ePTA and stabilized in LMP agarose inside the smallest possible tube relative to sample size. This maintained a vertical orientation of plant samples for optimal rotational axis during imaging. Low melting point agarose effectively coated and stabilized even the most delicate or complicated samples, and provided complete physical support preventing tissue movement during long scans. Higher scan magnifications and resolutions required longer scan times and our methods allowed both, which concomitantly enabled improved computational segmentation and analysis by providing higher quality raw image data.

Contrast enhancement with PTA has widespread utility due to broad staining of various plant tissues (Hayat, 1993; Staedler et al., 2013). As effective as PTA is as a contrast agent for most plant samples, it requires a lengthy incubation period—at least 14 days in our methods—to generate sufficient contrast. The application of microwave tissue processing could potentially accelerate PTA contrast enhancement protocols down to days or even hours (Kang et al., 1991; Zechmann and Zellnig, 2009; Carpentier et al., 2012). In our hands, conventional fixation in buffered aldehydes followed by post-fixation with osmium tetroxide and potassium ferrocyanide was effective to contrast samples like root tips and *Arabidopsis thaliana* plantlets, but penetration was limited into larger complex floral structures. Iodine-based contrast agents can rapidly contrast plant tissue within 24 hours, but they easily leach out of specimens into the surrounding agarose, lowering effective contrast and increasing scan noise and background.

Alternatively, more rigid plant specimens can be contrasted in iodine and instead stabilized with relatively X-ray transparent polyester fibers or small polystyrene beads, adding pieces of paper towel or lab wipes saturated with water or ethanol to maintain high relative humidity before sealing in sample tubes. Other robust plant material such as wood samples (Reynolds et al., 2018) or seeds and grains (Su and Xiao, 2020) are readily imaged with XRM with a minimum of preparation. Supplementary Figure 3 shows XRM scans of maize, sorghum, soybean, and wheat seeds where no fixation or contrast enhancement were used. Seeds were simply placed in PCR or Eppendorf tubes and stabilized with expanded polystyrene beads, to observe a range of internal seed structures.

Meristematic structures, which are often too large and dense for meaningful 3D imaging using fluorescence or EM, can greatly benefit from XRM. Numerous recent studies have provided important insights into meristematic and inflorescence structure development, including *Setaria viridis* (Yang et al., 2018), *Hordeum vulgare* (Walla et al., 2020), *Zea mays* (Teng et al., 2020), *Aquilegia coerulea* (Min and Kramer, 2020), and other grasses (Kellogg et al., 2013; AuBuchon-Elder et al., 2020). These examples illustrate the breadth of research questions that can be enhanced by additional cell-level 3D XRM imaging to measure and evaluate meristematic tissue development over scales not possible or practical with other methodologies. To address the challenge of meristematic cells having relatively uniform density, nuclei can be used as foci for segmentation since ePTA imparts high contrast to nucleic acids and the location of the nucleus in the cell can assist computational efforts to identify individual cells and cell walls.

Use of contrast agents for meristematic tissue is critical if cell-level segmentation is the goal. Thus, the development and application of cell wall-specific chemistries such as electron-dense probes or particles to generate XRM contrast agents that are specifically selective for plant and fungal cell walls would be highly beneficial. While selective staining may be possible using small molecules, we also envision the application of genetically encoded expression of ascorbate peroxidase (APEX2) fluorescent protein fusions to a library of select plant cellular compartments (Ariotti et al., 2018). This would allow osmium reactive diaminobenzidine deposits at target structures that could be visualized first by XRM and subsequently processed for high-resolution EM studies as described previously (Ng et al., 2016; Tsang et al., 2018; Kong et al., 2020).

Plant-microbe interactions, root biology, and vascular bundles were readily differentiated by XRM imaging with relatively straightforward sample preparation. For example, root nodules induced on *G. max* were visualized in detail (Figures 4A-D) without the time consuming and technically challenging tasks of physical sectioning, acquiring photomicrographs, and manually assembling images into a 3D model in order to study root nodule morphology and vasculature (Livingston et al., 2019). Mycorrhizal fungi that form arbuscules within cortical cells of plant roots for mutualistic nutrient exchange between plant and fungus can be visualized and quantified as shown by segmentation of *Rhizophagus irregularis* in maize roots (Figures 4E,F). We also developed a novel syringe barrel system where both host root system architecture and fungal symbiont were imaged *in situ* (Figures 4G,H, Supplementary Figure 4) without perturbation of the local environment. We are currently exploring how affinity-based probes with electron dense compounds or particles can be used to enhance fungal contrast for overall positional mapping as well as high-resolution imaging within the plant host (Scotson et al., 2019).

Cell-level measurement of roots in 3D is now possible with XRM imaging to better understand how simple root growth responses such as elongation, bending, and branching result in complicated and vast root networks that occupy square meters of soil (Figures 3A-D). This enables studies that compare cell number, morphology, and 3D position in root tips between *Z. mays* lines known to differ in root system architecture at relevant developmental stages (Jiang et al., 2019), or for modeling hydraulic conductivity (Heymans et al., 2020). Iodine-based contrast agents were useful for visualization of vascular elements (Figure 3E), and coupled with XRM will be of great value for mapping physically complex regions such as the root-shoot transition zone (nodal plexus) and graft junctions between rootstock and scion.

Imaging of EM prepared resin embedded specimens in the life sciences increasingly has been applied to 3D XRM (Hanssen et al., 2012). Likewise, protocol improvements in SBFEM sample preparation techniques (Wanner et al., 2015; Titze and Genoud, 2016) including the protocol used here with *en bloc* metal staining (Hua et al., 2015), demonstrated improved osmium penetration and enhanced contrast in thick plant tissues which inherently benefit high-resolution XRM and SBFEM correlative workflows (Bushong et al., 2015). While there have been a handful of published plant based SBFEM examples (Kittelmann et al., 2016; Oi et al., 2017; Czymmek et al., 2020; Harwood et al., 2020), we reasoned that further improvements in uniformity, conductivity and intensity of plant staining (Deerinck et al., 2018) would significantly benefit both XRM and SBFEM image contrast and quality for correlative studies. Additionally, specimens encapsulated in rigid hard formulation epoxy resins provided excellent stability for long duration XRM scans, translating to optimal resolution. Our protocol resulted in optically opaque specimens in resin blocks that initially were scanned with XRM to identify regions of interest and guided our subsequent high-resolution volume EM, all from the same block and using a single optimized sample preparation method.

Notably, we believe these data reflect the highest resolution reconstruction of subcellular plant structures via microCT or XRM published to date, allowing sub-organellar starch segmentation via deep learning (Fig 5 F,G). We envision that our plant-based correlative XRM to EM workflow will serve as an effective way to obtain 3D contextual histology, as a standalone approach, or modified to include other correlative strategies such as targeting genetically encoded structures and/or combined with light microscopy approaches. While less accessible, we acknowledge that further resolution gains using synchrotron-based platforms will allow nanoscale imaging (Hwu et al., 2017; Fonseca et al., 2018; Chin et al., 2020) also benefiting from ours and others’ plant sample preparation strategies. Ultimately, these tools and protocols will accelerate the ease in localizing, resolving and quantifying important discrete and multiscale plant biological phenomena, including developmental studies, phenotype characterization, and plant-microbe interactions, for subsequent high-resolution EM studies.

## METHODS

### Sample contrast and mounting

A preparation overview for specific samples is provided in Supplementary Table 1, as well as a workflow diagram illustrating how samples are fixed, stabilized, and mounted for XRM imaging in Supplementary Diagram 1. General methods and correlative microscopy details are described in the following section. Most samples were fixed directly in 20 mL glass vials in the contrast agent, ethanolic phosphotungstic acid (ePTA), using 1% PTA in 90% ethanol in ddH_2_O. A brief (~5 min) vacuum was applied to facilitate penetration of ePTA, particularly for hydrophobic plant tissue and floral structures with prolific trichomes, especially leaves. Samples remained in ePTA between 14 and 56 days on a mechanical lab rocker, with exchanges of ePTA every seven days. While incubation in ePTA less than 14 days gave inferior contrast and penetration, prolonged incubation did not adversely affect imaging and in most cases enhanced contrast. Some samples were fixed in 4% glutaraldehyde and 4% paraformaldehyde in 1X phosphate buffered saline (PBS) at pH 7.4. Samples were exposed to several cycles of vacuum/air until samples sunk after vacuum release. After aldehyde fixation overnight at 4°C, samples were rinsed 3 x 30 min in PBS, then post-fixed in 1% osmium tetroxide with 1% potassium ferrocyanide in PBS for 24 h at room temperature. Post-fixed samples were rinsed 3 x 60 min in PBS then stored at 4°C.

Fixed and contrast enhanced samples were rinsed 10 min in ddH_2_0 on a lab rocker. Low melting point (LMP) agarose (Promega #V2111) was prepared at 1% in ddH_2_O on a lab hot plate and removed from heat just as the agarose melted, prior to active boiling. A narrow transfer pipette was used to add enough agarose to the bottom of a 200 μm flat cap PCR tube to ensure the sample would be completely submerged when transferred. Fine forceps were used to transfer the sample from the ddH_2_O rinse vial, blotting briefly on a paper towel to remove excess ddH_2_O, then placed in the liquid agarose in the base of the tube. Additional agarose was added to completely fill the tube, avoiding air spaces or bubbles. After allowing agarose to set, the cap was closed tightly and a two-component epoxy gel (Devcon®, ITW) was used to completely seal the tube and to mount the tube on the end of a wooden applicator stick. Epoxy was allowed to harden for at least 60 min prior to loading into the XRM sample holder (process illustrated in Supplementary Diagram 1).

Low melting point agarose is our preferred medium for stabilizing small and delicate samples, particularly for long high resolution scans. If samples are larger and more robust, they can be stabilized in a variety of low density media, which then allow the use of iodine-based contrast agents such as Lugol’s iodine-potassium iodide solution. Lugol’s was prepared as 500 mL of a 4X stock solution by adding 50 g of potassium iodide to 400 mL ddH_2_O and stirring until dissolved, followed by adding 25 g iodine and adding ddH_2_O to a total volume of 500 mL. Polyester fibers typically used for pillows can be used to pack and cushion larger samples in appropriately sized plastic tubes. The fibers can be carefully moistened with ethanol or water—depending on whether the contrast agent was ethanolic or aqueous—before sealing the tube to ensure a stable relative humidity over the course of the scan. Similarly cotton wool can be used although the cotton fibers also absorb iodine (Lugol’s) which can complicate subsequent image segmentation. Finally, small expanded polystyrene beads in the 1mm-2mm range, available online or at most hobby/craft shops, can be used to stabilize larger samples like seeds, grains, and maize stalk segments.

### Correlative microscopy

*Nicotiana benthamiana* leaves were fixed, stained and embedded using a modified protocol for enhanced *en bloc* staining of large tissues (Hua et al., 2015) that was compatible for high resolution XRM and SBFEM imaging. Briefly, 3 mm biopsy punches of leaves were fixed overnight with 2% glutaraldehyde and 2% paraformaldehyde with 0.1% Tween in 0.1 M sodium cacodylate buffer, pH 7.4, rinsed in 3X in buffer and fixed for 2 h in 1% OsO_4_ in cacodylate buffer at room temperature. Samples were then transferred to 2.5% ferrocyanide in cacodylate buffer for 1.5 h, rinsed and placed in 1% thiocarbohydrazide (TCH, #21900; Electron Microscopy Sciences) in ddH_2_O for 45 min at 40°C. Subsequently, samples were then fixed with 2% OsO_4_ in ddH_2_O for 1.5 h, rinsed 2X in ddH_2_O and placed overnight in aqueous 1% uranyl acetate at 4°C. The samples were then transferred to an oven at 50°C for 2 h, rinsed, placed in lead aspartate at 50°C for 2 h, rinsed, dehydrated in a graded series of acetone and embedded in Embed 812 hard formulation epoxy resin.

Resin samples were trimmed, mounted with epoxy glue on an aluminum pin and imaged on a Zeiss Xradia 520 Versa using a 4X objective (Supplementary Table 2). For 40X imaging, a Zeiss Xradia 620 Versa was used, pixel size was 0.19 μm isotropic, exposure time is 27 seconds, and 3001 projections taken over 360 degrees. No filter was used with a 70 kV and 8.5 W power. Following scans, Optirecon 2.0 (Advanced Reconstruction Toolbox) deconvolution processing was applied. For SBFEM, sample was then remounted on a Gatan 3View aluminum pin with 2-part conductive silver epoxy (Circuit Works, CW2400), trimmed to minimize excess resin and sputter coated with AuPd before sectioning on a Zeiss GeminiSEM300 equipped with a Gatan 3View at 1.2 kV with focal charge compensation (Deerinck et al., 2018) using 8000 x 8000 pixel image size and 5 nm pixel size (x-y) and 50 nm slice thickness (z).

### X-ray microscopy

Apart from the 40X scan described above, all scans were conducted on a Zeiss Xradia 520 Versa, equipped with four objective lenses (0.4X,4X,20X,40X), as well as a 3072 x 1944 pixel flat panel detector with 75 μm pixel pitch. Scan parameters for all scans are listed in Supplementary Table 2. Zeiss Reconstructor software was used to automatically or manually reconstruct 3D volumes from 2D scan data, and ORS Dragonfly software (Montreal, Canada) was used for data integration, visualization, and animation of the scan data, and to export image data as 2D 16-bit TIFF stacks. Fly-through animations of 2D image stacks for scans shown in all Figures, as well as 3D volume rendering animations of selected scans, are available for download from (TBD).

### Image analysis and segmentation

Data from XRM scans were segmented using Amira software and with the assistance of a Wacom tablet for manual segmentation. Segmentation for Figure 1D used automatic methods as follows: 1) the inflorescence region was segmented using thresholding; 2) a distance map was generated from the inflorescence region; 3) H-maxima transform was applied to the distance map; 4) Watershed segmentation was used to segment individual branches. Segmentation for Figure 3B used automatic and manual methods as follows: 1) Based on thresholding, the mask image of root tip was created; 2) H-maxima transform was applied to the mask image; 3) watershed segmentation was applied to segment individual cells; 4) different cell classes were labeled manually.

Segmentation for Figure 3C used semi-automatic method as follows: 1) different thresholding values were used to segment different layers; 2) morphological operations (opening and closing) were applied to smooth the boundaries of layers; 3) boundaries were manually corrected or drawn through the cross-section images. The individual cells of the soy nodule in Figures 4C,D were segmented using marker based watershed segmentation as follows: 1) symbiosomes were segmented using a threshold value; 2) nuclei were segmented using a higher threshold value; 3) a marker based watershed method with nuclei as markers, was applied to segment individual cells. The vasculature was manually segmented from a TIFF stack of the volume from Figure 4B using Amira and a Wacom tablet and then combined with an automatic segmentation using region growing and morphological closing operation. Segmentation for Figure 4F used automatic and manual methods as follows: 1) Based on thresholding, the mask image of root was created; 2) H-maxima transform was applied to the mask image; 3) watershed segmentation was applied to segment individual cells; 4) Arbuscules were segmented using thresholding and component filtering. For Figure 4H, spores were manually segmented from the TIFF stack using Amira and a Wacom tablet. The original grayscale volume was then cropped along the Z-axis to show the locations of the spores throughout the volume. For correlative XRM data (Figure 5F), we used ORS Dragonfly Pro Deep Learning Module with the Sensor3D model (Novikov et al., 2019) which takes three slices into account for each slice segmented with Dice loss and AdaDelta optimization method for training. We then used a 128 x 128 pixel image patch with a stride ratio of 0.5 and a slice step of 2 (the slice 2 below and 2 above are used as input along with the current slice). Twenty slices were trained and data augmentation set to a factor of 15, with rotation, horizontal and vertical flipping, shearing (of 2 degrees) and scaling (0.9 to 1.1).

## ACKNOWLEDGEMENTS

Financial support for the X-ray microscope came from a research collaboration agreement between Valent BioSciences, Sumitomo Chemical Company, and the Donald Danforth Plant Science Center to C.N. Topp. Numerous colleagues kindly provided wild type plant material for development of imaging protocols outlined in this manuscript, in addition to advice and guidance on correctly describing plant structures in the Figures. Jiani Yang and Yunqing Yu (DDPSC) provided *Setaria viridis*, Dierdra Daniels and Daniela Floss (Valent BioSciences) provided *Rhizophagus irregularis* for *Z. mays* colonization and *Gigaspora margarita* for our syringe barrel culture system, Shrikaar Kambhampati (DDPSC) provided all soybean and pennycress floral material, Adam Bray and Dhineshkumar Thiruppathi (DDPSC) provided *Z. mays* stalk segments and nodal plexus samples, Penelope Lindsay (Cold Spring Harbor Labs) provided ear primordium samples of *Z. mays*, and David Weglarz (Still630) provided wheat kernels. Kind thanks to Tomo Kawashima (Univ. Kentucky) for additional plant morphology guidance for Figure 2, and Elizabeth (Toby) Kellogg, Blake Meyers, and Andrea Eveland (DDPSC), and Daniela Floss (Valent BioSciences) for helpful comments that greatly improved this manuscript. We acknowledge the Advanced Bioimaging Laboratory (RRID:SCR_018951) at the Donald Danforth Plant Science Center for support with correlative sample preparation including screening samples with LEO 912AB TEM acquired through an NSF Major Research Instrumentation grant (DBI-0116650). We acknowledge the DOE BER (DBI-0116650) to K. Czymmek, as well as Doug Allen and Kevin Chu (DDPSC) for providing tobacco plants for correlative work.

Finally, we thank Rachna Parwani, Robin White, Joel Mancuso and Ruth Redman (Zeiss) for excellent support collecting and processing the correlative high-resolution XRM and SBFSEM dataset.

## AUTHOR CONTRIBUTIONS

K.E.D., K.J.C., and C.N.T. designed the research; K.E.D. and K.J.C. performed the research; N.J., A.C.T., and K.J.C. provided computational analyses; K.E.D., K.J.C., and C.N.T. wrote the article.

## SUPPLEMENTARY MATERIAL

**Supplementary Diagram.**
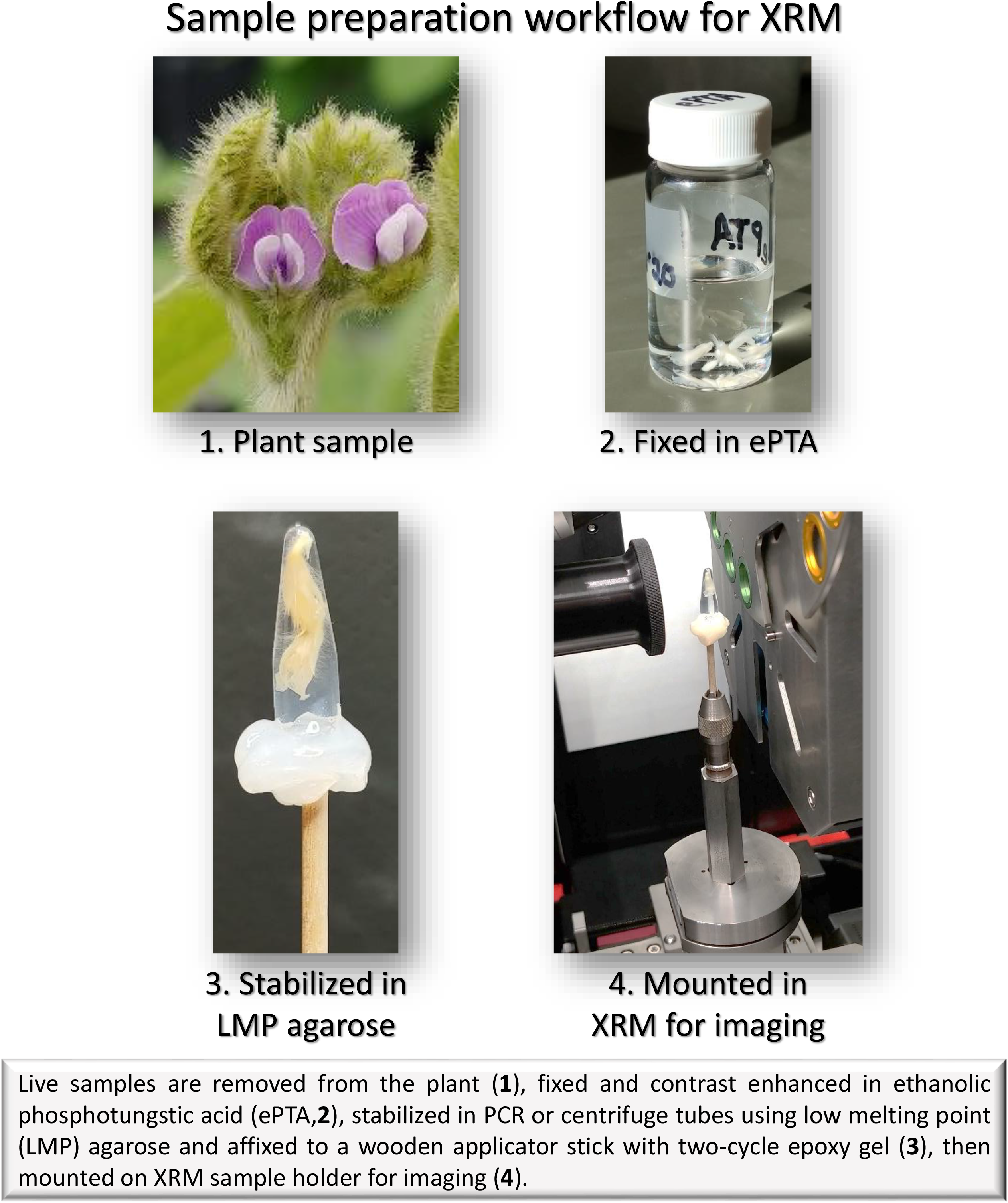
Sample preparation workflow for XRM Live samples are removed from the plant (**1**), fixed and contrast enhanced in ethanolic phosphotungstic acid (ePTA, **2**) stabilized in PCR or centrifuge tubes using low melting point (LMP) agarose and affixed to a wooden applicator stick with two-cycle epoxy gel (**3**), then mounted on XRM sample holder for imaging (**4**).

**Figure S1.**
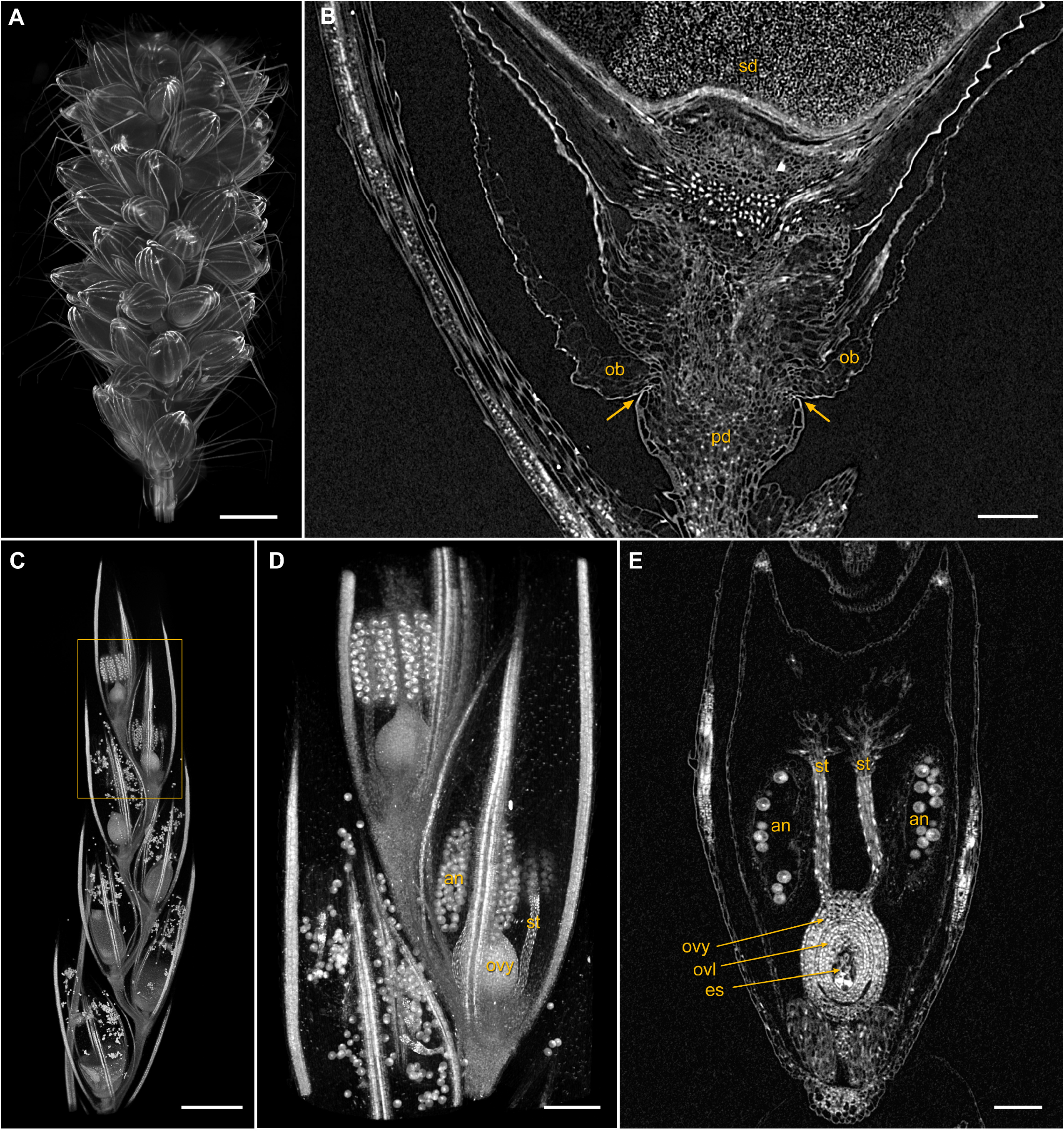
Inflorescence development **A** 3D volume rendering of a *Setaria viridis* inflorescence with seed-containing spikelets and sterile bristles visible; scale bar 3 mm. **B** High resolution image of a single spikelet from **A,** arrows indicate the region of the outer bracts **(ob)** and pedicel **(pd)** where the abscission zone will form as the seed **(sd)** matures; scale bar 100 μm. **C** 3D volume rendering of inflorescence structure from *Eragrostis tef(teff);* scale bar 1 mm. **D,E** High resolution scan of indicated region from **C.** Structures such as the ovary **(ovy),** anthers **(an),** and stigmas **(st)** are readily distinguished in the 3D volume rendering **(D),** with the ovule **(ovl)** and embryo sac **(es)** visible in the 2D clip plane **(E);** scale bars 200 μm **(D),** 100 um **(E),**

**Figure S2.**
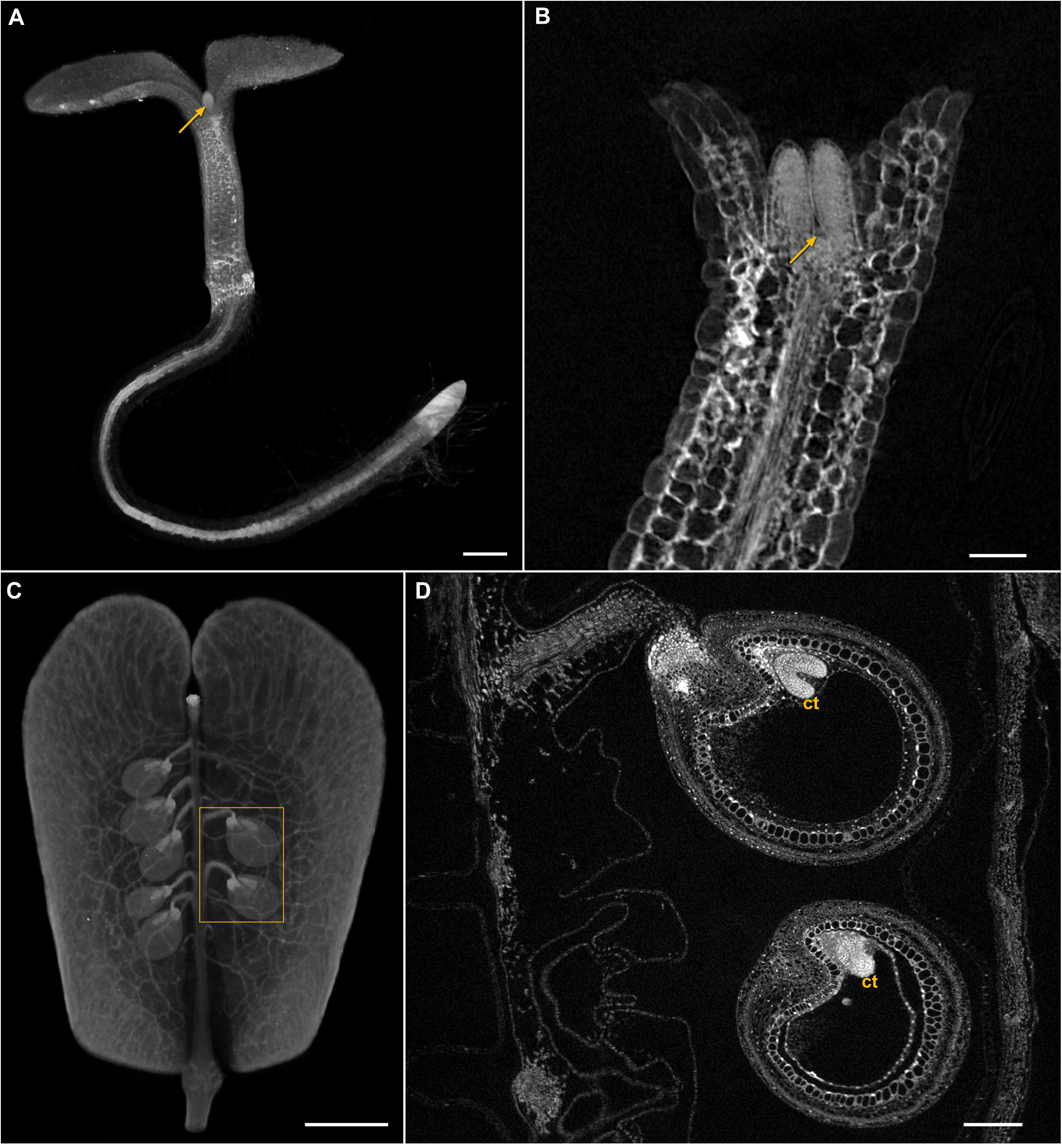
Multiscale imaging Examples of how the same sample can be imaged over multiple scales without removing it from the instrument. **A,B***Arabidopsis thaliana* seedling fixed and contrast enhanced, stabilized in low melting point **(LMP)** agarose, and imaged at low **(A)** and high **(B)** resolution. Arrows point to the developing shoot apical meristem; scale bars 200 um **(A),** 50 μm **(B).** Single pod at seed-filling stage of *Th/aspi arvense* (pennycress), fixed and contrast enhanced, stabilized in agarose, and imaged at low **(C)** and high **(D)** resolution. Cotyledons **(ct)** and multiple seed layers are clearly visualized; scale bars 1.5 mm **(C),** 200 μm **(D).**

**Figure S3.**
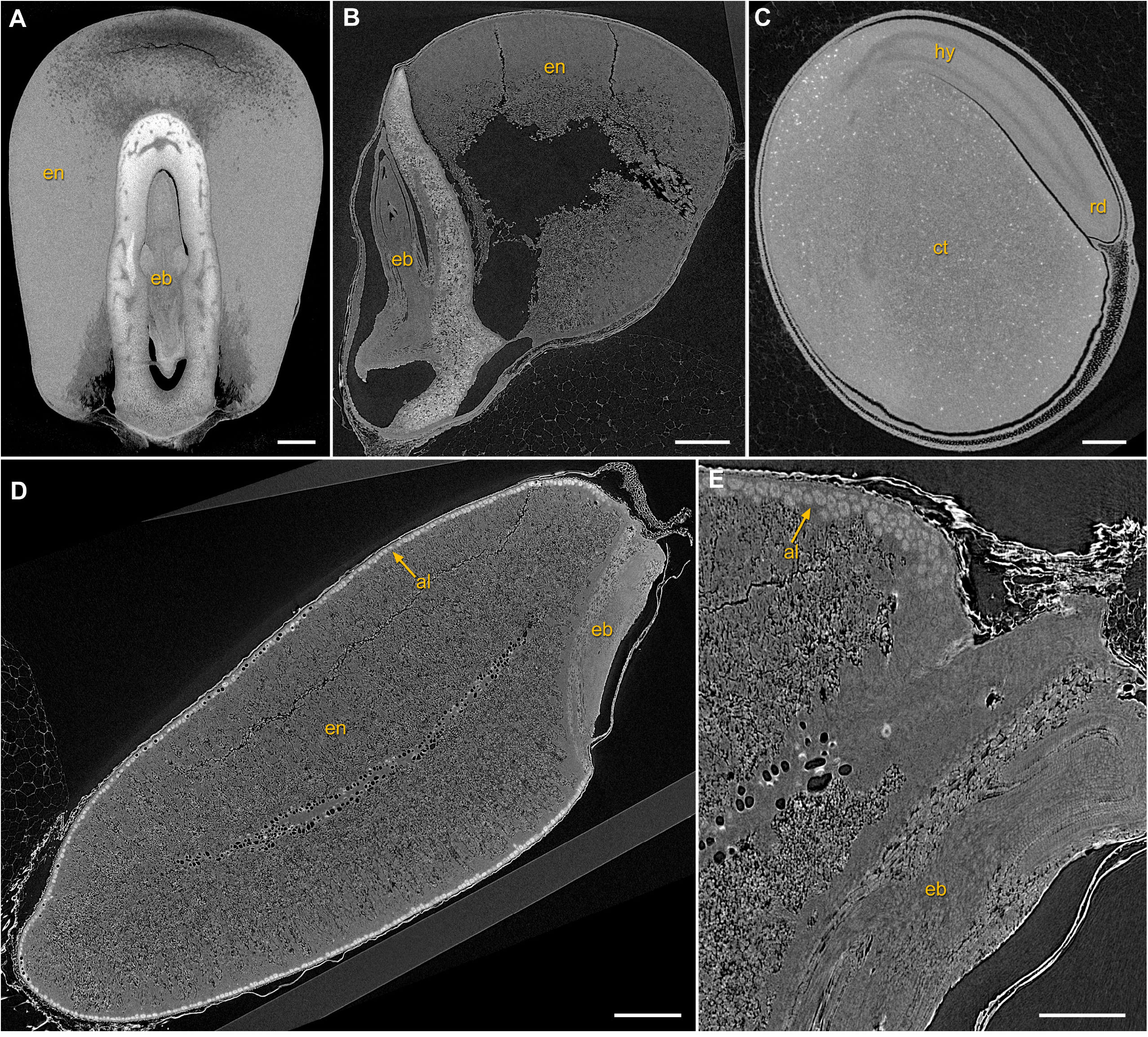
Seed morphology Seeds from maize **(A),** sorghum **(B),** soybean **(C),** and wheat **(D,E)** were scanned without fixation or contrast enhancement. Samples were stabilized in PCR or centrifuge tubes packed with expanded polystyrene beads. Internal structures such as endosperm **(en),** embryo **(eb),** cotyledon **(ct),** hypocotyl **(hy),** radicle **(rd),** and aleurone layer **(al)** are readily visualized; scale bars 1 mm **(A),** 500 μm **(B,C,D),** 200 μm **(E).**

**Figure S4.**
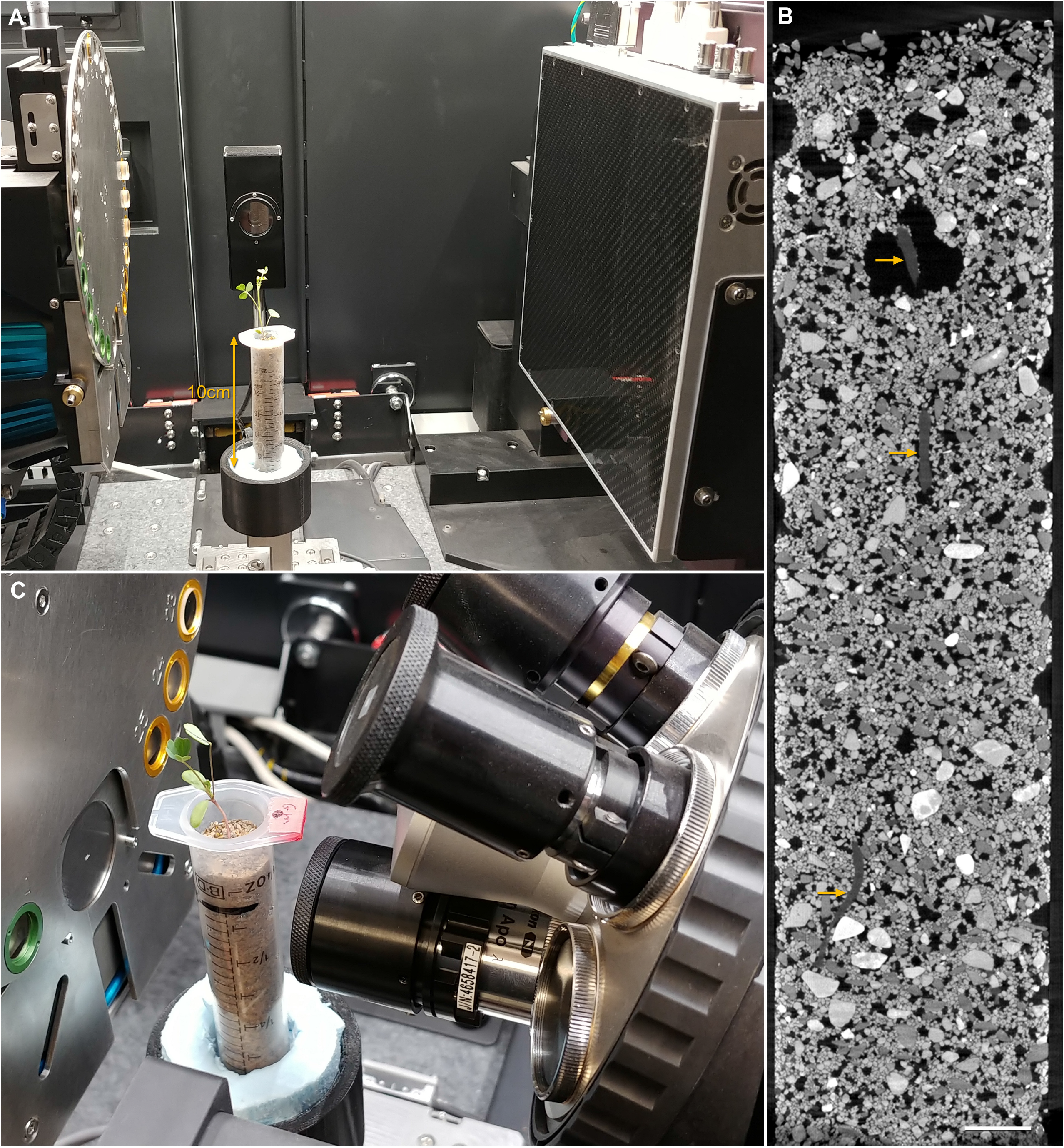
Multiscale *in situ* imaging of host-microbe interaction *Medicago sativa* (alfalfa) and mycorrhizal fungus *Gigaspora margarita* interaction visualized over a wide range of magnifications. Imaging with a flat panel detector **(A)** allows visualization of alfalfa root system architecture in the entire 2 cm x 10 cm syringe barrel volume **(B, arrows);** scale bar 5 mm. Image data from **B** directs high resolution 3D imaging with a 4X objective lens **(C),** where segmentation of individual G. *margarita* spores is presented in **Fig 4G,H.**

**Supplementary Table 1.**
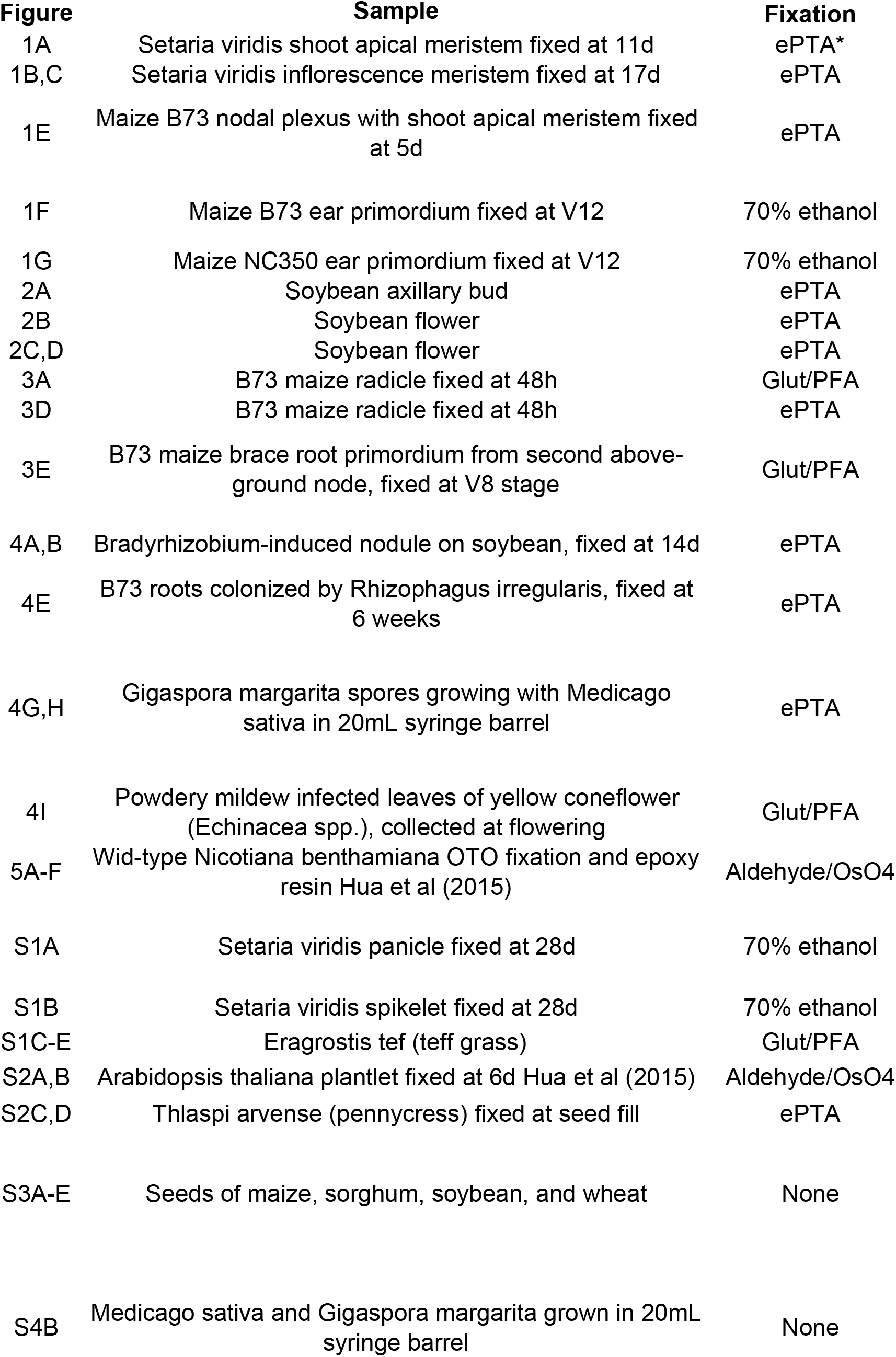
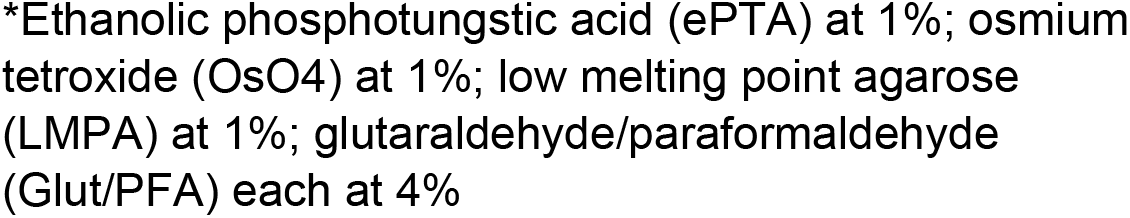
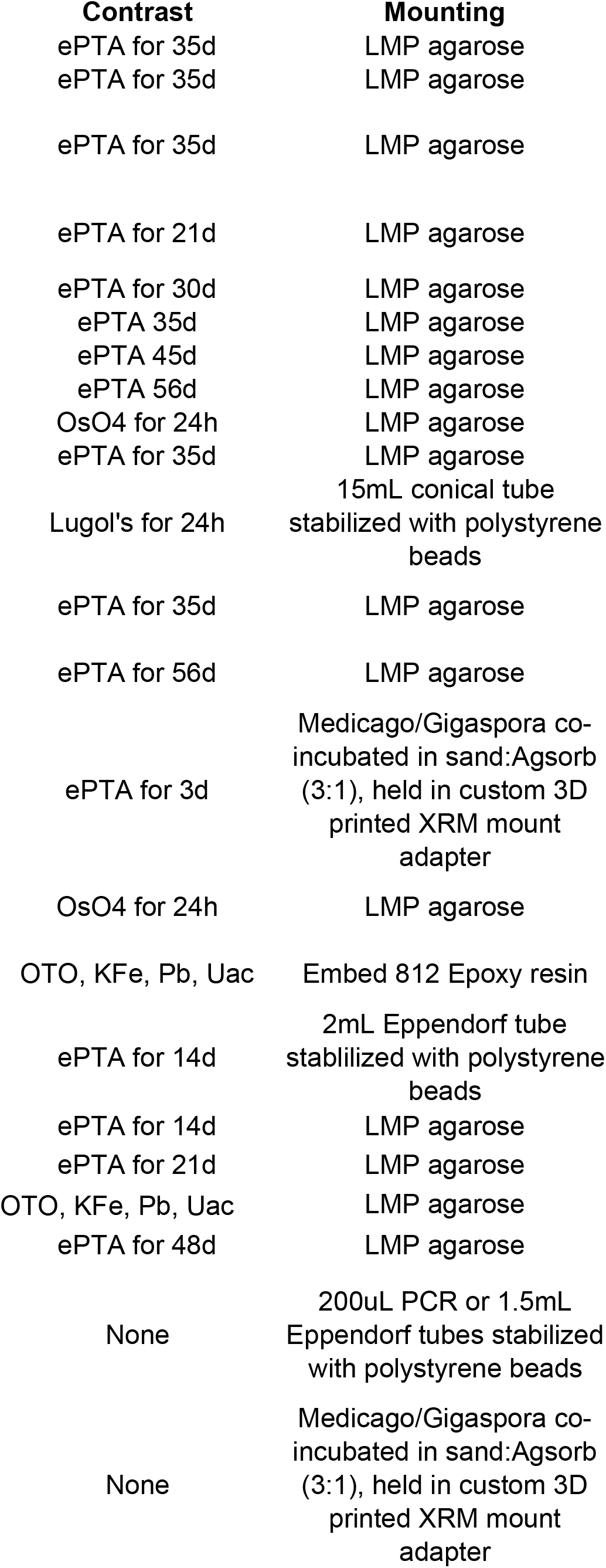
Sample preparation details for Figures and Supplementa

| **Figure** | **Sample** | **Fixation** |
| --- | --- | --- |
| 1A | *Setaria viridis* shoot apical meristem fixed at 11d | ePTA* |
| 1B,C | *Setaria viridis* inflorescence meristem fixed at 17d | ePTA |
| 1E | Maize B73 nodal plexus with shoot apical meristem fixed at 5d | ePTA |
| 1F | Maize B73 ear primordium fixed at V12 | 70% ethanol |
| 1G | Maize NC350 ear primordium fixed at V12 | 70% ethanol |
| 2A | Soybean axillary bud | ePTA |
| 2B | Soybean flower | ePTA |
| 2C,D | Soybean flower | ePTA |
| 3A | B73 maize radicle fixed at 48h | Glut/PFA |
| 3D | B73 maize radicle fixed at 48h | ePTA |
| 3E | B73 maize brace root primordium from second above-ground node, fixed at V8 stage | Glut/PFA |
| 4A,B | Bradyrhizobium-induced nodule on soybean, fixed at 14d | ePTA |
| 4E | B73 roots colonized by *Rhizophagus irregularis*, fixed at 6 weeks | ePTA |
| 4G,H | *Gigaspora margarita* spores growing with *Medicago sativa* in 20mL syringe barrel | ePTA |
| 4I | Powdery mildew infected leaves of yellow coneflower (*Echinacea* spp.), collected at flowering | Glut/PFA |
| 5A-F | Wid-type *Nicotiana benthamiana* OTO fixation and epoxy resin Hua et al (2015) | Aldehyde/OsO4 |
| S1A | *Setaria viridis* panicle fixed at 28d | 70% ethanol |
| S1B | *Setaria viridis* spikelet fixed at 28d | 70% ethanol |
| S1C-E | *Eragrostis tef* (teff grass) | Glut/PFA |
| S2A,B | *Arabidopsis thaliana* plantlet fixed at 6d Hua et al (2015) | Aldehyde/OsO4 |
| S2C,D | *Thlaspi arvense* (pennycress) fixed at seed fill | ePTA |
| S3A-E | Seeds of maize, sorghum, soybean, and wheat | None |
| S4B | *Medicago sativa* and *Gigaspora margarita* grown in 20mL syringe barrel | None |

\*Ethanollic phosphotungstic acid (ePTA) at 1%; osmium tetroxide ( $\text{OsO}_4$ ) at 1%; low melting point agarose (LMPA) at 1%; glutaraldehyde/paraformaldehyde (Glut/PFA) each at 4%

ry Figures
| **Contrast** | **Mounting** |
| --- | --- |
| ePTA for 35d | LMP agarose |
| ePTA for 35d | LMP agarose |
| ePTA for 35d | LMP agarose |
| ePTA for 21d | LMP agarose |
| ePTA for 30d | LMP agarose |
| ePTA 35d | LMP agarose |
| ePTA 45d | LMP agarose |
| ePTA 56d | LMP agarose |
| OsO4 for 24h | LMP agarose |
| ePTA for 35d | LMP agarose |
| Lugol's for 24h | 15mL conical tube<br>stabilized with polystyrene<br>beads |
| ePTA for 35d | LMP agarose |
| ePTA for 56d | LMP agarose |
| ePTA for 3d | Medicago/Gigaspora co-<br>incubated in sand:Agisorb<br>(3:1), held in custom 3D<br>printed XRM mount<br>adapter |
| OsO4 for 24h | LMP agarose |
| OTO, KFe, Pb, Uac | Embed 812 Epoxy resin |
| ePTA for 14d | 2mL Eppendorf tube<br>stabilized with polystyrene<br>beads |
| ePTA for 14d | LMP agarose |
| ePTA for 21d | LMP agarose |
| OTO, KFe, Pb, Uac | LMP agarose |
| ePTA for 48d | LMP agarose |
| None | 200uL PCR or 1.5mL<br>Eppendorf tubes stabilized<br>with polystyrene beads |
| None | Medicago/Gigaspora co-<br>incubated in sand:Agisorb<br>(3:1), held in custom 3D<br>printed XRM mount<br>adapter |

**Supplementary Table 2.** X-ray microscope scan parameters for imaging shown in Figures and Supplementary Figures.

## Footnotes

1 high-resolution image stacks (“flythroughs”) and videos portraying the 3D data sets can be found here: https://figshare.com/s/944efc8832e47fd4f203

